# MeCP2 Represses the Activity of Topoisomerase IIβ in Long Neuronal Genes

**DOI:** 10.1101/2023.02.24.529960

**Authors:** Sabin A. Nettles, Yoshiho Ikeuchi, Chibueze Agwu, Azad Bonni, Harrison W. Gabel

## Abstract

A unique signature of neuronal transcriptomes is the high expression of the longest genes in the genome (e.g. >100 kilobases). These genes encode proteins with essential functions in neuronal physiology, and disruption of long gene expression has been implicated in neurological disorders. DNA topoisomerases resolve topological constraints that arise on DNA and facilitate the expression of long genes in neurons. Conversely, methyl-CpG binding protein 2 (MeCP2), which is disrupted in Rett syndrome, can act as a transcriptional repressor to downregulate the expression of long genes. The molecular mechanisms underlying the regulation of long genes by these factors are not fully understood, however, and whether or not they directly influence each other is not known. Here, we identify a functional interaction between MeCP2 and Topoisomerase II-beta (TOP2β) in neurons. We show that MeCP2 and TOP2β physically interact *in vivo* and map protein sequences sufficient for their physical interaction *in vitro*. We profile TOP2β activity genome-wide in neurons and detect enrichment at regulatory regions and gene bodies of long neuronal genes, including long genes regulated by MeCP2. Further, we find that knockdown and overexpression of MeCP2 leads to altered TOP2β activity at MeCP2-regulated genes. Our findings uncover a mechanism by which MeCP2 inhibits the activity of TOP2β at long genes in neurons and suggest that this mechanism is disrupted in neurodevelopment disorders caused by mutation of MeCP2.

## Introduction

During development, diverse combinations of genes are expressed in order to establish the complex morphology, connectivity and function of neurons (Closser et al., 2022). The distinct complexity of the neuronal transcriptome is evidenced by the observation that the longest genes in the genome are expressed at uniquely high levels in the brain (Gabel et al., 2015). Many of these long pre-mRNAs (e.g. > 100 kilobases) encode proteins essential for neuronal function, including ion channels, synaptic receptors and cell adhesion molecules (Gabel et al., 2015), and have been implicated in neurological disorders (King et al., 2013). Long genes are expressed many fold higher in neurons than other cell types (Gabel et al., 2015; Sugino et al., 2014; Zylka et al., 2015) and analyses of transcript levels in developing mice and *in vitro*-derived human neurons demonstrate that the expression of long genes is a hallmark of functional maturity (McCoy et al., 2018). Pharmacological interventions that selectively downregulate long genes have been shown to lead to synaptic dysfunction and reduced neuronal transmission (Mabb et al., 2014). Thus, the expression of long genes is a distinctive and necessary feature of the neuronal transcriptome and aberrant expression of these genes represents a vulnerability in neurons that can lead to dysfunction.

Members of the topoisomerase gene family have been identified as essential for neural development and critical for the expression of long genes (Feng et al., 2017; King et al., 2013). Topoisomerases resolve DNA supercoiling and other topological constraints that arise during cellular processes including cell division, gene transcription and chromatin remodeling (Corbett and Berger, 2004; Nitiss, 2009; Roca, 2009). Type I topoisomerases (TOPI) relax DNA by transiently nicking and rejoining one strand of duplex DNA, whereas type II topoisomerases (TOP2) transiently break and rejoin both strands of duplex DNA simultaneously (Austin and Marsh, 1998; Koster et al., 2005; Pommier et al., 2016; Stewart et al., 1998). Mammalian cells express two genetically distinct TOP2 enzymes, TOP2*α* and TOP2β. Although TOP2*α* and TOP2β have similar structures and biochemical activities, they have different expression patterns and biological roles (Austin et al., 1993; Jenkins et al., 1992). For example, TOP2*α*is highly expressed in proliferating cells but excluded from neurons, whereas TOP2β is highly expressed in both dividing and post-mitotic cells (Capranico et al., 1992; Harkin et al., 2016; Juenke and Holden, 1993; Kondapi et al., 2004; Tiwari et al., 2012; Tsutsui et al., 1993; Watanabe et al., 1994). Consistent with this expression pattern, TOP2*α* is essential for cell-cycle-related events, such as DNA replication, whereas TOP2β plays an essential role in DNA repair, transcription, and development. Germline or conditional deletion of *Top2b* leads to defective brain development and perinatal death (Lyu and Wang, 2003; Yang, 2000). Pharmacological inhibition or knockdown of TOP2β in cultured neurons also leads to reduced expression of long genes in neurons (King et al., 2013). While the exact mechanism by which TOP2β promotes gene expression is not clear, TOP2β may decatenate DNA to open chromatin and activate regulatory regions (Ju et al., 2006; Lyu et al., 2006) or unwind DNA to aid the progression of RNA polymerase though genes (King et al., 2013; Mabb et al., 2014; Zylka et al., 2015). Together, these findings indicate that TOP2βis critical for the robust expression of long genes, which are essential for normal brain function.

Balanced gene regulation is essential for neuronal function and recent studies have implicated methyl-CpG-binding protein 2 (MECP2) as an important regulator of transcription in neurons that preferentially represses long genes (Boxer et al., 2020; Chen et al., 2015; Gabel et al., 2015; Lagger et al., 2017; Renthal et al., 2018; Sugino et al., 2014). Loss or overexpression of MeCP2 causes the severe neurological disorders Rett syndrome (Amir et al., 1999) and MeCP2 duplication syndrome (van Esch et al., 2005), respectively, demonstrating its importance for nervous system function. Although multiple functions of MeCP2 have been described, molecular and genomic analyses support a model in which MeCP2 binds to methylated DNA within and around genes to promote a repressive chromatin structure and down-regulate gene transcription. Transcriptional repression by MeCP2 appears to be mediated in part by downregulation of enhancers within genes, leading to downregulation of transcription initiation (Clemens et al., 2020; Boxer et al., 2020; Chahrour et al., 2008; Lewis et al., 1992; Lyst et al., 2013; Nan et al., 1993a). Notably, because long genes are enriched for DNA methylation and contain many enhancers, these repressive effects most robustly affect long genes (Clemens et al., 2020). Consistent with the essential roles for long genes in neurons, MeCP2 represses genes that are critical for neuronal development and physiology (Gabel et al., 2015; Sugino et al., 2014). These studies suggest that an important function of MeCP2 is to preferentially tune down the expression of long genes and implicate disruption of long gene regulation in the pathology of MeCP2-associated neurodevelopmental disorders.

Mechanistically, biochemical studies of MeCP2 have identified protein interactors that associate with MeCP2 to mediate its functions. Extensive evidence demonstrates MeCP2 may function as a repressor by interacting with the NCoR/SMRT-corepressor complex (Lyst et al., 2013). In addition, interactors such as PSIP1/LEDGF and TCF-20 may also play important roles in MeCP2-mediated gene regulation (Jian et al., 2022; Li et al., 2016). However, whether additional protein interactors or mechanisms play a key role in MeCP2 functions, remains to be investigated.

In this study, we identify a physical interaction between MeCP2 and TOP2β with important functional implications for the regulation of long genes in neurons. Through unbiased proteomic analysis and targeted interaction mapping, we identify and interrogate the physical association between TOP2β and MeCP2, implicating a key domain of MeCP2 required for this interaction. We profile the sites of TOP2βactivity in neurons genome-wide and demonstrate that TOP2β is preferentially active at long genes, including genes repressed by MeCP2. We further demonstrate that altering MeCP2 levels in neurons alters TOP2β activity at these long, MeCP2-regulated genes. Altogether, our findings demonstrate an interaction between TOP2β and MeCP2 in the regulation of essential neuronal genes, that when disrupted, may contribute to MeCP2-related neurological disorders.

## Results

### MeCP2 selectively binds TOP2β in the mouse brain

To explore the molecular mechanism underlying MeCP2 transcriptional regulation of long genes, we sought to identify additional proteins that may interact with MeCP2 in neurons. We expressed and purified FLAG-tagged MeCP2 protein in cultured cortical neurons and performed unbiased mass spectrometry analysis of co-precipitating proteins (**Table S1**). In addition to previously identified interactors of MeCP2 (KPNA3, PSIP1/LEDGF), TOP2β was reproducibly identified in this analysis (**Figure 1A**). Motivated by previous findings from separate studies showing that inactivation of topoisomerase enzymes (King et al., 2013) and mutation of MeCP2 (Gabel et al., 2015) leads to opposing effects on long gene expression, we sought to further investigate this interaction. To independently test the association of TOP2β-MeCP2 and assess its functional relevance in the postnatal brain, we immunoprecipitated MeCP2 from forebrain extracts of wild-type 8-week mice. To eliminate the potential for nucleic acid bridging, nuclear extracts were treated with benzonase, a DNA and RNA nuclease, before the immunoprecipitation (IP) was carried out. Through this endogenous co-immunoprecipitation (Co-IP) analysis, we found MeCP2 interacts with TOP2β but not TOP1, the other major topoisomerase that is robustly expressed in the brain (**Figure 1B**). Together, the detection and validation of this interaction between MeCP2 and TOP2β suggest that modulation of long gene expression by MeCP2 may occur through a mechanism that involves the activity of TOP2β.

**Figure 1.**
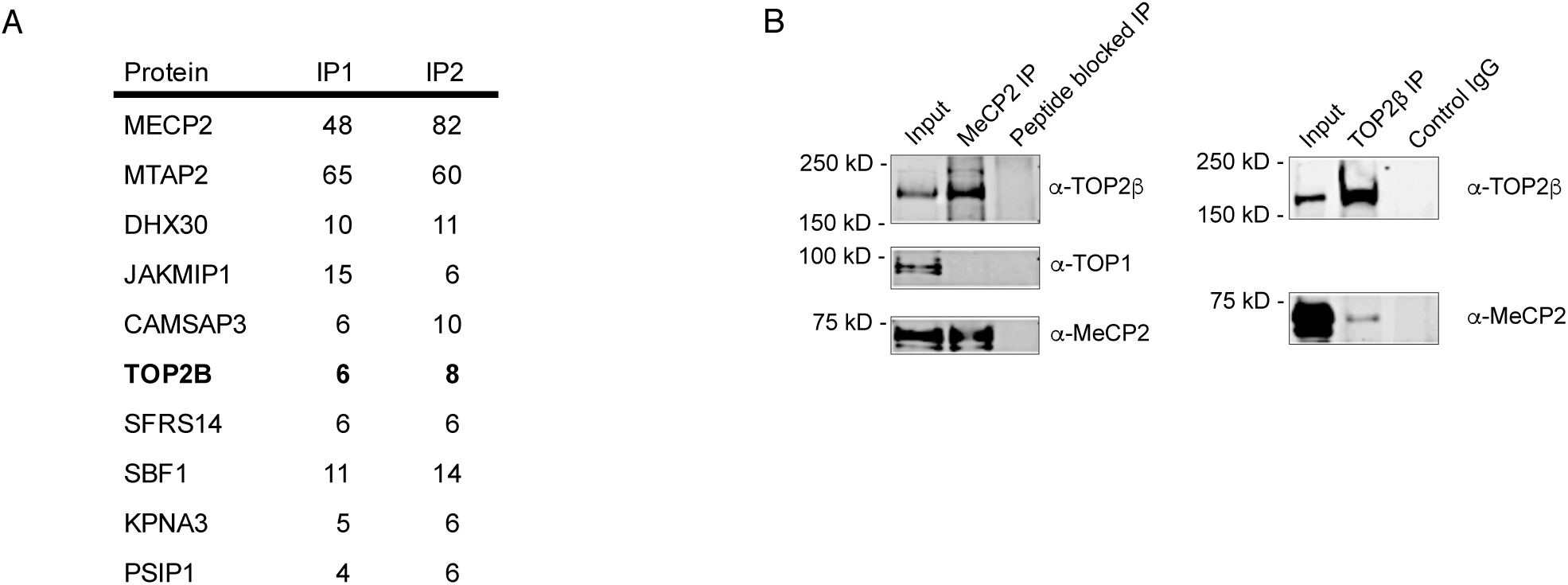
**MeCP2 interacts with TOP2β in neurons.** (A) MeCP2 was isolated by FLAG immunoprecipitation from lysates of mouse cortical neurons infected with lentivirus expressing FLAG-MeCP2 from the CMV (IP1) or Synapsin1 (IP2) promoter. Interacting proteins were detected by liquid chromatography with tandem mass spectrometry (LC/MS/MS) and quantified by CompPASS analysis. The ten proteins with the highest average weighted D scores are shown for proteins detected in the two LC/MS/MS analyses (see Table S1). Total spectral count for each protein is shown for each analysis. (B) Left, immunoprecipitation (IP) of MeCP2 from nuclear extracts from the whole cortex of wild-type mice, using an antibody against MeCP2 shows co-IP of TOP2βbut not TOP1. Peptide blocked IP indicates IP performed with antibody against MeCP2 that was pre-incubated with the peptide to which the antibody was raised. MeCP2-bound endogenous proteins were assayed on western blots. Right, IP of TOP2β with co-IP of MeCP2 from neuronal nuclear extracts from whole cortex of wild-type mice. Control IgG indicates IP performed with an antibody against another protein, CtBP.

The proteins MeCP2 and TOP2βare highly conserved across vertebrate species and are well-characterized, with specific domains having been defined as important for distinct protein functions (Austin et al., 1995; Baker et al., 2013; Caron and Wang, 1994; Heckman et al., 2014a; Lewis et al., 1992; Lyst et al., 2013; Nan et al., 1993a; Wang, 1996; Watt and Hickson, 1994). Thus, determining sequences in each protein required for the MeCP2-TOP2βassociation might provide insight into the nature and importance of the protein-protein interaction. We therefore established an *in vitro* co-IP assay in heterologous cells and asked which sequences within each protein are sufficient for the interaction. Co-expression of MYC-tagged full length (1-1612aa) TOP2β with FLAG-tagged full length (1-484aa) MeCP2 in HEK293 cells led to a detectable interaction upon co-IP (**Figure 2A**). Notably, this interaction was specific under these conditions as expression of MYC-tagged full length TOP2β with another nuclear protein, FLAG-tagged full length (1-908aa) DNMT3A, in these cells did not lead to a detectable co-IP signal. This validated the interaction we detected *in vivo* and allowed us to begin to dissect the amino acids that mediate the association between MeCP2 and TOP2β.

**Figure 2.**
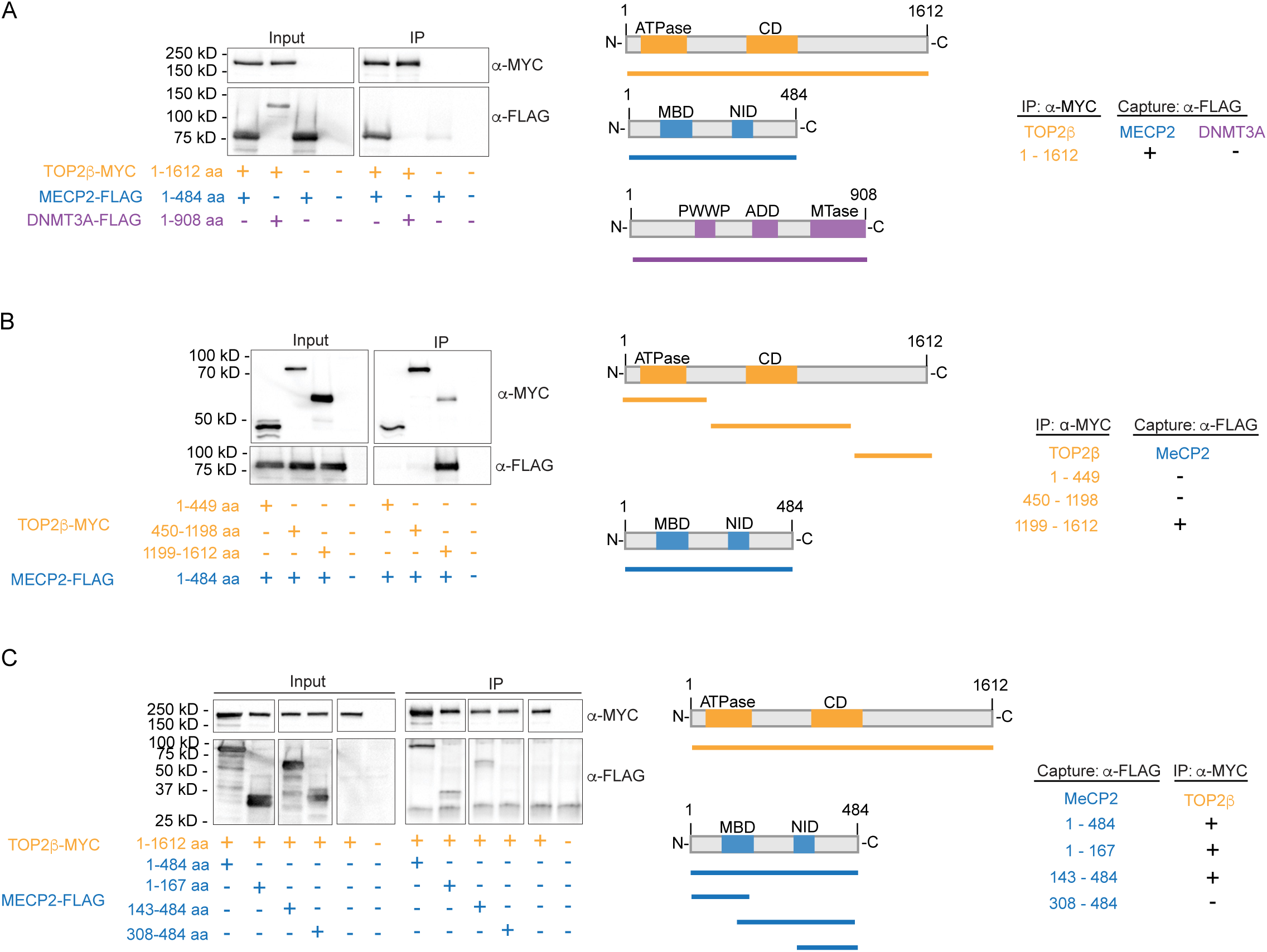
**Identification of protein regions sufficient for the MeCP2-TOP2β interaction.** (A) Co-immunoprecipitation analysis of MYC-tagged full-length TOP2β, FLAG-tagged full-length MeCP2 and FLAG-tagged full-length DNMT3A in HEK293T cells. Immunoprecipitation of full-length TOP2β binds full-length MeCP2 but failed to bind another nuclear protein, DNMT3A. Left, cell lysates were immunoprecipitated with an antibody against MYC, and co-precipitated FLAG was visualized by western blots. Center, overview of TOP2β (orange), MeCP2 (blue) and DNMT3A (purple) domain structure. Right, summary of results. Domains are annotated as follows: CD, catalytic domain containing the active tyrosine; MBD, methyl-DNA-binding domain; NID, NCoR-interaction domain; PWWP, proline-tryptophan-tryptophan-proline domain; ADD, auto-inhibitory ATRX-DNMT3A-DNMT3L domain; MTase, methyltransferase domain. (B) Co-immunoprecipitation analysis of MYC-tagged fragments of TOP2β with full-length FLAG-tagged MeCP2 in HEK293T cells. Full-length MeCP2 interacts with the C-terminal region (1199-1612) of TOP2βand, but not with the N-terminal region (1-499) or the central region (450-1198). Left, cell lysates were immunoprecipitated with an antibody against MYC and co-precipitated FLAG was visualized by western blot. Lysates from untransfected cells were run in the last lane of the blot as a control. Center, overview of TOP2β (orange) and MeCP2 (blue) domain structure. Right, summary of results. (C) Co-immunoprecipitation analysis of FLAG-tagged fragments of MeCP2 with MYC-tagged full-length TOP2β. IP of full-length TOP2β recovers full-length (1-484) MeCP2, a fragment spanning the majority of the MBD (1-167), and a fragment spanning part of the MBD and the NID (143-484) but fails to detect a fragment containing the C-terminal region (308-484) of MeCP2. Left, cell lysates were immunoprecipitated with an antibody against MYC, and co-precipitated FLAG was visualized by western blot. Lysates from untransfected cells were run in the last lane of the blot as a control. Center, overview of TOP2β (orange) and MeCP2 (blue) domain structure. Blue lines show deletion fragments of MeCP2. Right, summary of results. All blots in this figure are representative of at least two biological replicate experiments HEK293T cell transfections.

We next probed the sequences within TOP2β that are sufficient for the interaction with MeCP2. TOP2β contains three functional regions. The amino-terminal region contains an ATP-binding domain that allows dimerization with other TOP2 monomers. The central region of TOP2β is the catalytic domain which contains the active tyrosine responsible for the strand breakage and re-ligation. The carboxyl-terminal region is suggested to be required for localization, regulation of enzyme activity by post-translational modification, and regulation of enzyme function by protein-protein interactions. (Austin and Marsh, 1998; Austin et al., 1993, 1995; Berger et al., 1996; Chang et al., 2013; Chen et al., 2013; Jenkins et al., 1992; Lindsley and Wang, 1991). To determine the region of TOP2β that binds MeCP2, we co-expressed exogenous MYC-tagged N-terminal (1-449aa), central catalytic (450-1198aa), and C-terminal (1199-1612aa) fragments with FLAG-tagged full length MeCP2 (1-484aa) in HEK293 cells. This analysis detected co-IP of MeCP2 specifically with the C-terminal region of TOP2β, whereas the N-terminal region and the catalytic region of TOP2β do not interact with MeCP2 (**Figure 2B**). The interaction between the MeCP2 and the C-terminal region of TOP2β is notable because the N-terminal ATPase and central breakage/reunion domains are very similar between TOP2*α* and TOP2β, whereas the C-terminal regions differ both in size and sequence and is considered the domain that gives rise to the enzyme-specific functions of TOP2α and TOP2β (Gilroy and Austin, 2011; Linka et al., 2007a; Meczes et al., 2008).

We next interrogated the region of MeCP2 required for the interaction with TOP2β. MeCP2 contains two critical domains: the methyl-DNA-binding domain (MBD), which is the region of MeCP2 that binds methylated DNA, and the NCoR-interaction domain (NID), required for the association with the NCoR corepressor complex (Guo et al., 2014; Heckman et al., 2014b; Lewis et al., 1992; Lister et al., 2013; Lyst et al., 2013; Nan et al., 1993a). Collectively, these two domains contribute to the repressive role of MeCP2 during transcription. A minimal protein containing the two domains is sufficient to rescue the multiple effects of MeCP2 loss in mice that model the severe neurological disorder Rett syndrome, which is caused by MeCP2 mutation (Tillotson et al., 2017). To determine the region of MeCP2 that is required for interaction with TOP2β, we co-expressed a FLAG-tagged MeCP2 fragment that contains the MBD (1-167aa), a fragment spanning the MBD and NID (143-484aa), and a C-terminal fragment (384-484aa), together with MYC-tagged full length TOP2β (1-1612aa). The fragments containing the MBD of MeCP2, 1-167aa and 147-484aa, were efficiently precipitated by MYC-tagged full length TOP2β, while the C-terminus of MeCP2 did not interact (**Figure 2C**). These results suggest that TOP2β interacts with the MBD of MeCP2. Notably, this is one of the key domains for rescue of the Rett phenotype, raising the possibility that loss of TOP2β interaction could contribute to Rett syndrome.

### TOP2β activity is enriched within long genes in neurons

Previous analyses have shown that pharmacological inhibition or knockdown of TOP2β in neurons causes reduced expression of long genes, but the site of TOP2β action during the transcription of long genes is not well understood. We therefore sought to identify where TOP2β acts in neurons in order to understand how this activity could be modulated by MeCP2. To gain insight into where on the genome TOP2β is active in neurons, we performed etoposide-mediated topoisomerase-immunoprecipitation (eTIP-seq) (Sano et al., 2008) in primary neurons isolated from mouse embryonic day 14.5 (E14.5) cerebral cortex. In this genomic profiling assay, etoposide treatment of cells is used to link topoisomerase to the DNA and profile its activity across the genome (**Figure 3A**). Etoposide is an inhibitor of TOP2 proteins that traps the enzyme in a covalently linked complex with the cleaved DNA at the intermediate step in its cutting and re-ligation cycle (Osheroff, 1989; Wu et al., 2011). Thus, in contrast to a conventional chromatin immunoprecipitation (ChIP) assay, which uses formaldehyde to non-specifically crosslink protein to DNA and profile binding in the genome, employing eTIP allowed us to directly assess the sites of TOP2β activity in neurons and study how altering MeCP2 impacts TOP2β activity.

**Figure 3.**
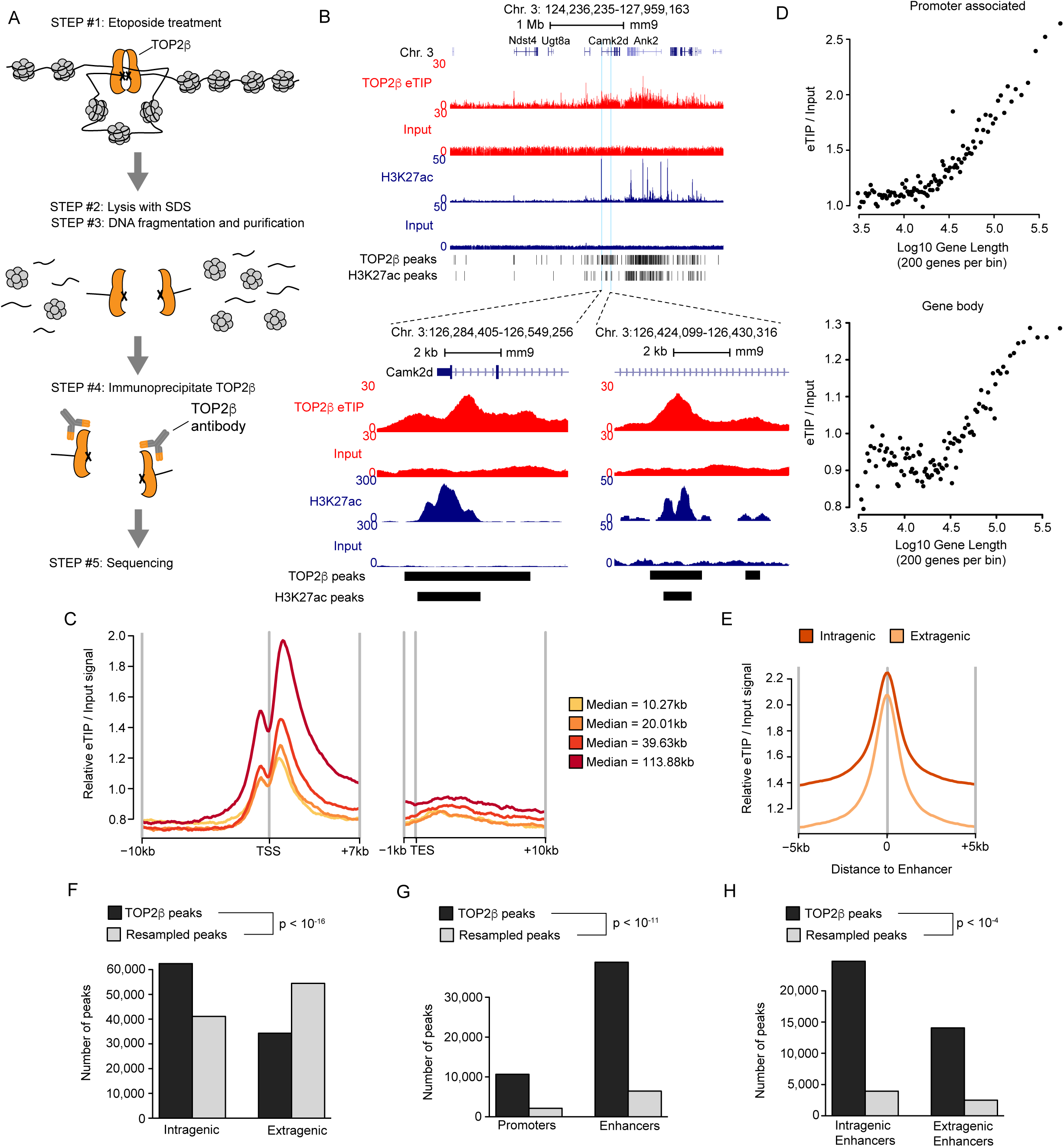
**TOP2β is preferentially active at long genes in neurons.** (A) Schematic representation of the eTIP-seq procedure to directly map sites of TOP2β activity, adapted from (Sano et al., 2008). TOP2β cleaves a DNA duplex to make a gate for a second DNA duplex to pass through. In the eTIP assay, etoposide treatment freezes TOP2β in its cleavage complex, leaving it covalently linked to its site of action. Following denaturing and fragmentation under stringent conditions containing SDS, TOP2β-linked DNA is immunoprecipitated, purified and sequenced. (B) Genome browser snapshots of eTIP-seq and H3K27ac ChIP-seq from DIV12 cultured cortical neurons. Top, an ∼2Mb region of the genome displaying the profile for TOP2β eTIP, H3K27ac and their respective inputs. Enrichment of eTIP-seq signal is more prominent at long genes (e.g. *Camk2d*) compared to shorter genes (e.g. *Ugt8a*). Bottom left, zoomed in view of the promoter region of the *Camk2d* gene. Bottom right, zoomed in view of an enhancer region of the *Camk2d* gene. Peaks of TOP2β eTIP and H3K27ac called by the MACS2 algorithm are indicated below the tracks. Gene annotations and scales are depicted above. (C) Aggregate plot of input-normalized eTIP signals at genes divided into quantiles of gene length (see methods). The median gene length for each group is indicated. Average signal around the transcription start site (TSS) and transcription end site (TES) is shown. Mean values plotted for 100bp bins. (D) Running average plots of input-normalized eTIP signal at the promoter associated region (top) and gene bodies (bottom), from eTIP-seq. Averages are shown for bins of 200 genes ranked according to gene length. (E) Aggregate plot of input-normalized eTIP signal at putative intragenic and extragenic enhancers genome-wide in cultured cortical neurons. Putative enhancers were defined by H3K27ac peaks that did not overlap the TSS of a gene. The plot is centered at the midpoint for each enhancer and the surrounding 10kb region. Mean values plotted for 100bp bins. (F) Bar plots of the genomic distribution of TOP2β peaks and resampled TOP2β peaks at intragenic and extragenic regions. Resampled peaks were generated by shuffling TOP2β peak-sized regions randomly around the genome. A Chi-squared test was conducted between the frequencies of TOP2β peaks and resampled TOP2β peaks and resulting p-values are indicated. (G) Bar plots of the genomic distribution of TOP2β peaks and resampled TOP2β peaks at promoters and enhancers. Resampling and statistical analysis were performed as described in panel F. (H) Bar plots of the genomic distribution of TOP2β peaks and resampled TOP2β peaks at intragenic enhancers and extragenic enhancers. Resampling and statistical analysis were performed as described in panel F.

To assess TOP2β activity in neurons, we utilized cultured cortical neurons at day *in vitro* 12 (DIV12) where we could administer etoposide directly to the cells and perform eTIP-seq (**Figure 3A**). Isolation of DNA by eTIP from etoposide treated cultured neurons yielded high levels of immunoprecipitated DNA compared to vehicle (DMSO) treated controls, indicating the high specificity of the assay (**Figure S1A, Table S5**). Deep sequencing of eTIP-seq samples revealed enrichment of TOP2β activity compared to input controls at local sites as well as broad regions (**Figure 3B**). Visual inspection of the eTIP profile relative to genome annotations showed enrichment of TOP2β at promoters and gene bodies. Notably, the eTIP signal appeared to be higher at longer genes (>100kb) relative to shorter genes (**Figure 3B**). To quantify TOP2β activity at all genes in the genome, we performed aggregate analysis and quantification of the signal at the promoters and gene bodies. Aggregate plots of promoter regions showed TOP2β is consistently associated with these regions, with the peak of the signal located just downstream of the transcription start site (TSS), spanning the TSS +1kb to +3kb region (hereafter referred to as “promoter associated” eTIP signal) (**Figure 3C**). Aggregate plots of genes binned by gene length revealed a length-associated enrichment of TOP2β at promoter associated regions and gene bodies of long genes, with the highest TOP2β signal at the longest genes and lower signal at the shorter genes (**Figure 3C**). Quantitative analysis of TOP2β signal at the promoter associated regions and gene bodies of all genes supported the aggregate profiles, showing a robust correlation between TOP2β enrichment and gene length (**Figure 3D, Table S2, S3**).

Long genes are robustly expressed in neurons, raising the possibility that TOP2β may be associated with these genes due to high expression in these cells. Therefore, we assessed whether the TOP2β signal at genes is correlated with gene expression, and if so, whether this association can explain the enrichment of TOP2βwith long genes. We performed RNA-seq analysis on neuronal cultures at an identical timepoint as eTIP-seq and assessed TOP2β enrichment as a function of gene expression using aggregate plots, in which genes were binned by level of gene expression (**Figure S1B**). This analysis showed a modest association between expression level and enrichment of TOP2β at promoters and within genes. Notably, however, while long genes are robustly expressed in our cultured neurons, we find that long genes are not more highly expressed than shorter genes in these cultures (**Figure S1C, Table S3**). This analysis indicates that TOP2β activity is associated with mRNA levels, but that recruitment of TOP2βto long genes cannot be explained by high expression of these genes alone.

We noted that additional sites of local eTIP enrichment can be observed outside of gene promoter regions and these sites are reminiscent of peaks associated with enhancers (**Figure 3B**). We therefore sought to assess the locations of TOP2β relative to these distal regulatory regions across the genome. To define active regulatory regions in the genome, we performed ChIP-seq on the histone modification Histone H3 lysine 27 acetylation (H3K27ac), a mark associated with active promoters and enhancers. We then performed peak-calling on these H3K27ac chip data to define active promoters and putative enhancer elements and assessed the degree of overlap of eTIP signal with these sites (**Table S4**). Examination of eTIP profiles revealed clear enrichment of TOP2β activity at acetyl peaks corresponding to putative enhancer elements (**Figure 3B**). Aggregate plot analysis of eTIP at these putative enhancer elements revealed enriched TOP2βsignal for enhancers found inside and outside of genes (**Figure 3E, Table S4**). Notably, enrichment of TOP2β was higher at enhancers located within genes (intragenic) and their surrounding regions relative to enhancers located outside of genes (extragenic) and their surrounding regions. This increased enrichment at intragenic enhancers may in part explain the increased signal inside long genes which can contain many enhancers.

To independently assess the profile of TOP2β activity genome-wide in an unbiased manner, we performed peak-calling on the TOP2βeTIP-seq signal. We then performed overlap analysis assessing where these peaks land in the genome, and compared them to resampled controls to assess if the distribution of TOP2β peaks differs from chance distributions. This analysis revealed that TOP2β peaks are significantly more prevalent within intragenic regions than extragenic regions compared to resampled controls (**Figure 3F, Table S4**). Analysis of TOP2β peak overlap with promoters and putative enhancers, as defined by H3K27ac peak calling analysis described above, indicated TOP2βpeaks fall within promoters and enhancers significantly more than by chance (**Figure 3G, Table S4**). We next assessed whether TOP2β peaks are more likely to fall within intragenic enhancers or extragenic enhancers. Our analysis shows TOP2β peaks that overlap with enhancers are biased toward associating with intragenic enhancers compared to resampled controls (**Figure 3H, Table S4**). Together this peak distribution analysis is consistent with TOP2β enrichment within genes, including at intragenic enhancers, as described above (**Figure 3E, Table S4**). Our analysis of the distribution of TOP2β peaks revealed additional TOP2β peaks in unannotated regions (**Figure S1D, Table S4**). Initial evaluation of these unannotated peaks suggests many of these sites may indeed be enhancers that were just below the level of detection of the H3K27ac peak calling analysis, however, many may not be enhancers. Most notable is the observation that nearly two-thirds of the unannotated TOP2β peaks are intragenic, suggesting TOP2β is engaged and active at many sites within these genes.

To further interrogate the profile of TOP2β activity across the genome and assess the validity of our findings, we performed eTIP-seq using a second, independently generated TOP2β antibody that targets a different region of the protein. Consistent with the initial observations, the second antibody showed TOP2βenrichment at promoter associated regions and gene bodies of long genes, as well as at enhancer regions **(Figure S2A – S2D, Table S4**). Peaks detected genome-wide were nearly three times more likely to fall within an intragenic region compared to an extragenic region (**Figure S2E, Table S4**), three times more likely to fall within an enhancer compared to a promoter (**Figure S2F, Table S4**), and two and a half times more likely fall within an intragenic enhancer compared to an extragenic enhancer (**Figure S2G, Table S4**). Furthermore, additional TOP2β peaks were observed in unannotated regions (**Figure S2H, Table S4**). Taken together, these analyses confirm TOP2β enrichment at promoters and enhancers within long genes, suggesting that the activity of TOP2β at these regulatory regions may play an important role in the transcription of these genes.

### MeCP2 regulates TOP2β activity at long genes repressed by MeCP2

The opposing effects of MeCP2 and TOP2β on long gene expression, along with our findings that MeCP2 and TOP2β interact in cells, raises the possibility that MeCP2 may inhibit TOP2β activity to affect transcription of long genes. We therefore sought to assess if TOP2β is active at MeCP2-regulated long genes and determine if MeCP2 modulates the activity of TOP2βat these genes.

Analysis of eTIP-seq signal at “MeCP2-repressed” genes, a set of long genes that have previously been shown to be consistently upregulated in expression when MeCP2 levels are low and downregulated in expression when MeCP2 levels are high across studies of MeCP2 mutations (Gabel et al., 2015), revealed TOP2β enrichment at the promoter associated regions and gene bodies of these genes. This signal was enriched relative to unchanged genes and “MeCP2-activated genes”, a set of genes that are downregulated in expression when MeCP2 levels are low and upregulated in expression when MeCP2 levels are high (**Figure 4A**). Quantitative analysis of eTIP signal at MeCP2-regulated genes show TOP2β is significantly enriched at the promoter associated regions and genes bodies of MeCP2-repressed genes compared to all other genes (**Figure 4B, Table S3**). In this analysis, we also find that the MeCP2-activated genes displayed some TOP2β enrichment relative to unchanged genes, albeit, not to the levels of the MeCP2-repressed genes. This effect may be related to the relative expression level of this population of genes in our system. Altogether, these observations are consistent with our findings of a length-associated TOP2β signature and indicate that MeCP2-repressed genes, like other long genes, are targets of TOP2β activity.

**Figure 4.**
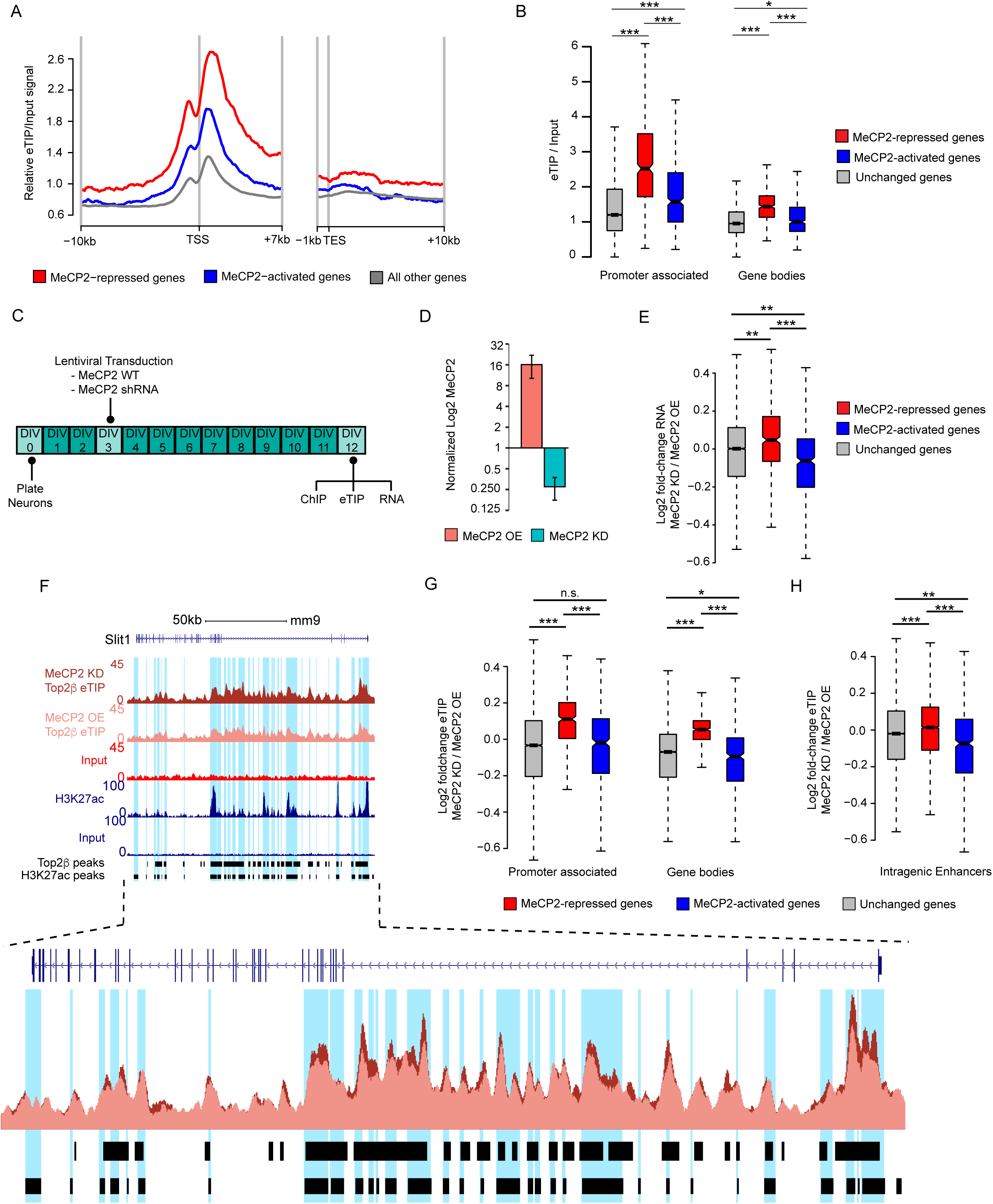
**Altering MeCP2 levels in neurons impacts TOP2β activity at MeCP2-regulated genes.** (A) Aggregate plot of input-normalized eTIP signal in DIV12 primary culture cortical neurons at MeCP2-repressed, MeCP2-activated, and all other genes. Gene sets are those previously identified as consistently dysregulated across multiple MeCP2 mutant datasets and brain regions (Gabel et al., 2015). (B) Boxplot of input-normalized eTIP signal at the promoter associated regions (left) and gene bodies (right) of MeCP2-repressed, MeCP2-activated, and all other genes. *, p < 0.01; ***, p < 10^-15^ Wilcoxon rank-sum test. (C) Schematic depicting experimental design to manipulate MeCP2 levels in primary culture cortical neurons. Embryonic neurons were isolated and cultured for 3 days before being transduced with exogenous wild-type MeCP2 (MeCP2 OE) or MeCP2 shRNA (MeCP2 KD) lentivirus. At DIV12, cells were harvested for downstream high-throughput genomic experiments (e.g. eTIP-seq, RNA-seq and ChIP-seq). (D) Relative MeCP2 RNA expression in DIV12 cortical primary neurons transduced with MeCP2 OE or MeCP2 KD lentivirus, as determined by RT-qPCR. n = 3 biological replicates per group. Graph shows *Actb* normalized mean ± SEM of MeCP2 expression relative to MeCP2 expression in control neurons in which MeCP2 expression was manipulated. (E) Boxplot of fold-changes in exonic RNA for MeCP2-repressed, MeCP2-actived and all other genes in DIV12 cortical neurons transduced with either MeCP2 OE or MeCP2 KD lentivirus. **, p < 10^-9^; ***, p < 10^-16^ Wilcoxon rank-sum test. (F) Top, genome browser snapshot of eTIP-seq signal at an example MeCP2-repressed gene, *Slit1*, in MeCP2 OE and MeCP2 KD DIV12 primary culture cortical neurons. Regions corresponding to H3K27ac peaks (putative enhancers and promoters) are highlighted in blue. Bottom, superimposition of eTIP-seq tracks, illustrates higher signal at this gene in the MeCP2 KD condition compared to the MeCP2 OE condition. (G) Boxplot of fold-changes in the promoter associated regions (left) and gene bodies (right) eTIP signal of MeCP2-repressed, MeCP2-actived and all other genes in DIV12 cortical neurons transduced with either MeCP2 OE or MeCP2 KD lentivirus. Fold-changes were calculated by edgeR analysis of eTIP signal at promoter associated regions and gene bodies. n.s., not significant; *, p < 0.05; ***, p < 10^-16^ Wilcoxon rank-sum test. (H) Boxplot of fold-changes in eTIP signal in DIV12 cortical primary neurons transduced with either MeCP2 OE or MeCP2 KD lentivirus for enhancers located within MeCP2-repressed, MeCP2-actived and all other genes. Fold-changes were calculated by edgeR analysis of eTIP signal at enhancer regions. **, p < 10^-9^; ***, p < 10^-16^ Wilcoxon rank-sum test.

Given that MeCP2 preferentially down-regulates a subset of neuronal long genes (e.g. MeCP2-repressed genes), we hypothesized TOP2β that is recruited to long genes may be negatively regulated by MeCP2, via the TOP2β-MeCP2 protein interaction, to restrict TOP2βactivity and transcriptional activation of these genes. We therefore asked whether manipulating MeCP2 expression would alter TOP2β activity across the genome. To address this question, we carried out knockdown (KD) or overexpression (OE) of MeCP2 in our established neuronal culture system and performed eTIP-seq (**Figure 4C, Table S3**). Because the level of MeCP2 protein increases dramatically over postnatal development (Balmer et al., 2003; Skene et al., 2010), our culture system contains low levels of MeCP2 compared to the adult brain. Thus, we overexpressed MeCP2 in these neuronal cultures to model the role of MeCP2 in the mature brain, while still providing access to the cells in order to perform eTIP-seq. We paired this with knockdown of MeCP2 to remove low levels of MeCP2 in the cultures and better recapitulate Rett syndrome-like conditions in which there is little to no MeCP2 activity present in neurons. We directly assessed MeCP2 RNA levels in the MeCP2 KD and MeCP2 OE cultured neurons using quantitative real-time PCR (RT-qPCR) (**Figure 4D, Table S5**). In the MeCP2 KD neurons, there was a robust reduction of endogenous MeCP2, while in the MeCP2 OE neurons, there was significantly higher MeCP2 expression.

To assess if our culture system mirrors aspects of the MeCP2-mediated gene regulation we observe in the brain, we performed total RNA-seq on the MeCP2 KD and MeCP2 OE cultured neurons in order to determine if the MeCP2-regulated genes identified in the *in vivo* studies of multiple brain tissues (Gabel et al., 2015) were misregulated in the same direction in the cultured neurons. We performed differential gene expression analysis and find that the MeCP2-repressed genes were significantly upregulated and the MeCP2-activated were significantly downregulated in MeCP2 KD neurons (**Figure 4E, Table S3**). This analysis suggests that our manipulation of MeCP2 in the cultured neurons elicits changes in gene expression that are consistent with the changes in gene expression observed in the brain, and therefore provides a suitable context to assess how levels of MeCP2 in neurons may affect TOP2β activity.

To investigate how MeCP2 expression alters TOP2β activity, we performed eTIP-seq in the MeCP2 KD and MeCP2 OE cultured neurons. Given that we observe TOP2β enrichment at the promoter associated regions and gene bodies of MeCP2-repressed genes, we hypothesized that loss of MeCP2 in the cultured neurons would induce an increase in TOP2β activity. Visualization of the eTIP signal in the MeCP2 KD and MeCP2 OE neurons at long, MeCP2-repressed genes suggested higher TOP2β activity in the MeCP2 KD neurons compared to MeCP2 OE neurons (**Figure 4F, Table S3**). Differential analysis of eTIP signal from MeCP2 KD and MeCP2 OE neurons revealed increased TOP2β activity at the promoter associated regions and gene bodies of MeCP2-repressed genes in the MeCP2 KD neurons (**Figure 4G, Table S3**). At MeCP2-acitvated genes we find the TOP2βactivity at promoter associated regions were unchanged while the gene bodies showed reduced signal in the MeCP2 KD neurons compared to MeCP2 OE neurons.

We previously showed one mechanism by which MeCP2 regulates long gene expression is by repressing the activity of intragenic enhancers associated with these genes (Clemens et al., 2020). To explore if TOP2β activity at these sites is affected by MeCP2, we evaluated the change in eTIP signal at intragenic enhancers of the MeCP2-regulated genes (Gabel et al., 2015) in the MeCP2 KD neurons compared to the MeCP2 OE neurons. Our analysis shows TOP2βactivity is subtly, but significantly, increased at MeCP2-repressed enhancers (**Figure 4H, Table S4**) when MeCP2 protein expression is low compared to when it is high. Together, these findings raise the possibility that negative regulation of TOP2β by MeCP2 may contribute to the repression of intragenic enhancers.

## Discussion

In this study, we have established a physical interaction in neurons between MeCP2 and TOP2β and demonstrated its importance for modulating TOP2β activity at long genes that are transcriptionally regulated by MeCP2. One model consistent with our findings is that when neurons are born in the developing brain, TOP2βis recruited to long genes and facilitates their expression. During postnatal development as neurons mature, MeCP2 becomes highly expressed and associates with methylated DNA in genes. At long, highly methylated genes, MeCP2 interacts with TOP2β to act as a molecular break on TOP2β activity within gene bodies and at enhancers, thereby leading to preferential restriction of TOP2β and tempering of gene expression. In the absence of MeCP2, as in Rett syndrome, the molecular break on TOP2β has been removed thereby allowing for TOP2βactivity to become unrestricted within long, MeCP2-regulated genes (**Figure 5**). This regulation may be key to the transcriptional regulatory role of MeCP2 to tune neuronal gene expression programs.

**Figure 5.**
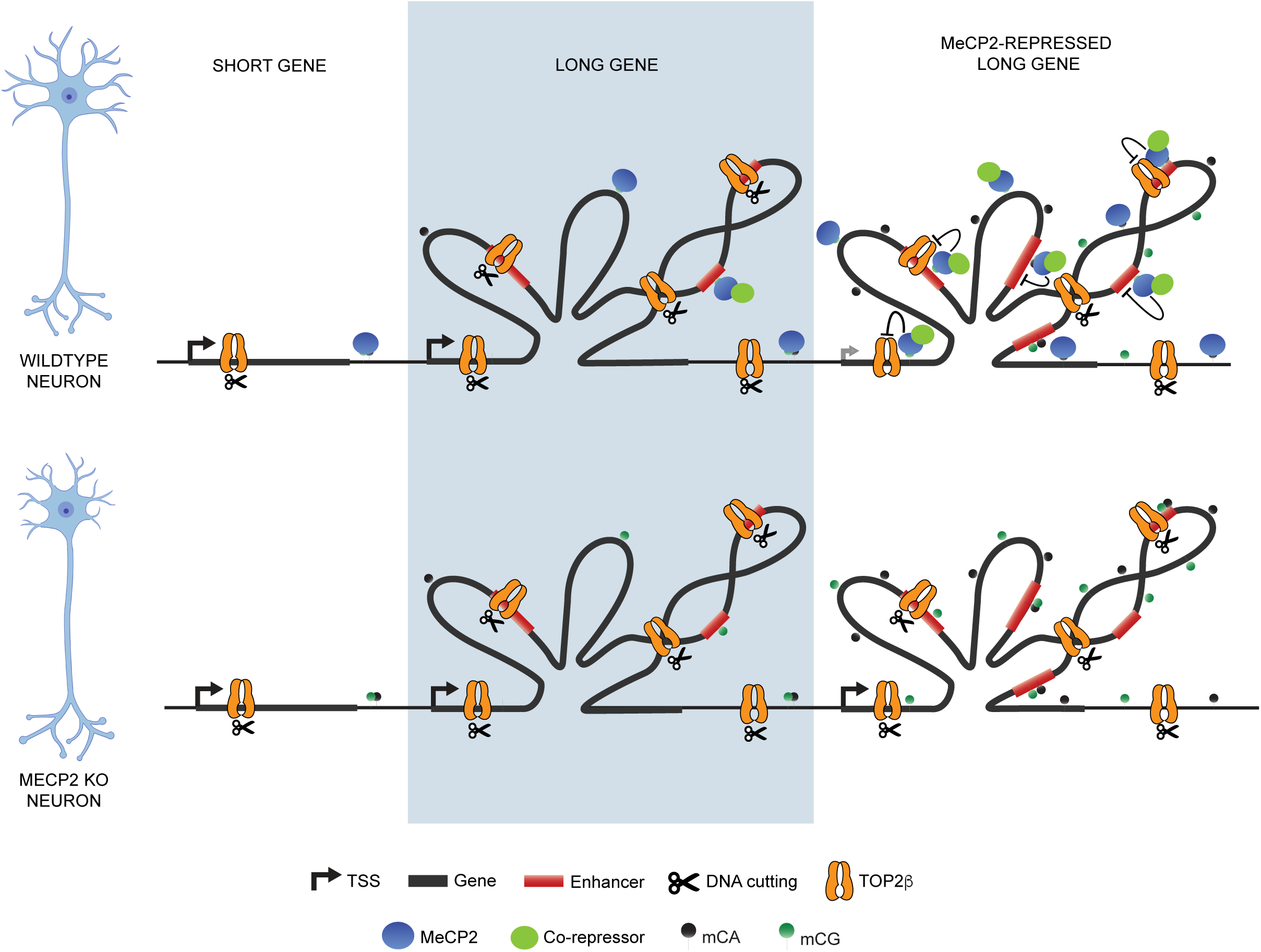
**A model of repression of TOP2β activity in neurons by MeCP2.** In neurons, TOP2β is preferentially recruited to promoter associated regions, gene bodies and enhancers of long genes, where it resolves topological constraints and facilitates gene expression. At a subset of long genes (“MeCP2-repressed genes”), DNA methylation (mCG and mCA dinucleotides) and MeCP2 binding are enriched. MeCP2 interacts with TOP2β, acting as a molecular break on activity of TOP2β to fine-tune the expression of these genes. When MeCP2 is absent (MeCP2 KO) the molecular break on TOP2βis removed, causing overactivity of TOP2βand overexpression of these long, MeCP2-repressed genes.

Type I and type II topoisomerases have been shown to promote the expression of long genes critical for neuronal development and synapses by cutting the DNA and resolving topological constraints. Through biochemical analysis, we show MeCP2 binds uniquely to the type II topoisomerase, TOP2β, but not the type I topoisomerase, TOP1, both of which are highly expressed in neurons. This is notable because TOP2 enzymes have been shown to have extended C-terminal sequences that play important roles in regulating these enzymes. Using fragments to map the regions sufficient for the interaction between MeCP2 and TOP2β, we show MeCP2 interacts with the C-terminal region (CTR) of TOP2β. Structural and functional studies have revealed that the CTR is the most divergent region between of TOP2α and TOP2β, and that removal of the CTRs of TOP2α and TOP2β did not affect the overall catalytic activity of either enzyme (Austin and Marsh, 1998; Linka et al., 2007b). However, truncation of the TOP2β CTR caused increased binding of TOP2β to DNA, whereas truncation of the TOP2α CTR had no effect on the binding of TOP2α to DNA, suggesting the CTR of TOP2β may have a negative regulatory function (Gilroy and Austin, 2011; Meczes et al., 2008). Collectively, these observations provide evidence of the CTR as a potentially important regulatory site of TOP2βactivity, particularly in neurons. This specific interaction may have evolved in the vertebrate lineage, where the expression and specialization of TOP2β, TOP2α, and MeCP2 have emerged (Austin & Marsh, 1998, de Mendoza et al., 2021), and may contribute to the regulation of complex vertebrate neuronal transcriptomes.

Our mapping of fragments of MeCP2 sufficient for the interaction with TOP2β implicates this interaction in Rett syndrome-related biology. Our study shows the TOP2β interaction occurs for MeCP2 fragments that contain the C-terminal portion of the MBD. In structural analysis, this portion of the MBD is found outside of the region that directly contacts the DNA (Nan et al., 1993b). This suggests that MeCP2 can interact with TOP2β and regulate its activity while bound to DNA. Previous studies have highlighted two critically important domains of MeCP2 for proper function, the MBD and the NID (Lyst et al., 2013; Nan et al., 1993a; Tillotson et al., 2017). The MBD domain of MeCP2 is required for the interaction with methylated DNA (Gabel et al., 2015; Guo et al., 2014; Lagger et al., 2017; Lister et al., 2013), while the NID domain of MeCP2 is required for the interaction with the NCoR-corepressor complex in cells (Lyst et al., 2013). Notably, our findings suggest the need to fully evaluate the effects of Rett syndrome causing missense mutations, which have been shown to be concentrated in the MBD, as some mutations may in fact disrupt the interaction with TOP2β rather than solely disrupting methyl-DNA binding capacity.

While the mechanism by which TOP2β facilitates long gene expression is not known, our study identifies TOP2β sites of action within long genes that may provide insight into its function in neurons. By using eTIP-seq to profile TOP2β activity across the genome, we specifically recovered TOP2β enzymes in the process of cutting double-stranded DNA, thereby associating functional relevance to each site of enrichment. Previous studies in primary mouse cortical neurons and mouse cerebellar neurons observed TOP2β enrichment by ChIP-seq at actively transcribed genes, promoters, across regions of open chromatin, and adjacent to genes that encode ion channels and receptors (Madabhushi et al., 2015; Sano et al., 2008; Tiwari et al., 2012). To our knowledge, we are the first to report a genome-wide gene-length-associated enrichment of TOP2β activity in neurons. Globally, we find TOP2β is distributed throughout the gene body with TOP2β activity markedly higher in long genes relative to shorter genes. Interestingly, we also observe significant TOP2β enrichment just downstream of promoters and at intragenic enhancers. Future studies can assess how TOP2β activity is targeted specifically to longer genes, and what role it plays in facilitating the expression of these genes.

MeCP2 has been characterized as a repressor given its ability to bind methylated DNA and repress transcription via the recruitment of the NCoR-corepressor complex. Loss or disruption of MeCP2 has been shown to cause dysregulation of intragenic enhancers and up-regulation of long, highly-methylated genes which may contribute to Rett syndrome pathology (Boxer et al., 2020; Clemens et al., 2020). Our results provide evidence that TOP2β plays a role in MeCP2-mediated repression of long genes. By profiling TOP2βgenome-wide in neurons that have low initial levels of MeCP2, we have identified TOP2β enrichment at the promoter associated regions and gene bodies of long, MeCP2-repressed genes (Gabel et al., 2015). This demonstrates that, like other long genes, MeCP2-repressed genes are preferential targets of TOP2β. Manipulating MeCP2 expression in cultured neurons to levels similar to adult wild-type (MeCP2 OE) or adult knockout (MeCP2 KD) alters the activity of TOP2β at MeCP2 regulated genes in a manner that is consistent with MeCP2 inhibiting TOP2β at these sites. Thus, our findings provide insight into the functional significance of the interaction between MeCP2 and TOP2βin the regulation of long genes. Because MeCP2 does not possess a catalytic domain, it is considered to function as a bridge between chromatin and the NCoR co-repressor complex (Lyst and Bird, 2015). One possible model for this function is that MeCP2, through association with the NCoR complex, inhibits TOP2β activity to repress the expression of long genes. The NCoR complex contains a histone deacetylase (HDAC3) and may function to deacetylate TOP2β. Future studies can investigate whether post-translational modifications are altered on TOP2β in MeCP2 mutants.

Outside of a direct role in long gene expression, our findings, combined with evidence from recent studies, may also implicate MeCP2-TOP2β regulation as an important modulator of DNA damage of these genes in neurons. TOP2βcuts DNA to facilitate changes in DNA topology and if TOP2β activity is disrupted during this process, it can lead to an aborted cutting and ligation cycle that result in DNA damage in the form of a double-stranded break (DSB). Because long genes experience topological constraints that do not otherwise affect shorter genes, long genes may require higher levels of TOP2βactivity to resolve these constraints. In the context of high TOP2β activity at long genes, MeCP2 may be an important modulator to down-regulate TOP2β activity within optimal levels. When MeCP2 is absent, TOP2β activity may be overactive, giving rise to additional errors and increased DSBs. Notably, previous studies have reported recurrent DSBs in long genes in neural progenitors (Wei et al., 2016) and detected increased levels of DSBs in neural progenitors from of *Mecp2* knockout mice (Alessio et al., 2018). Furthermore, a recent genetic suppressor screen identifying mutations in genes that ameliorate Rett-like phenotypes in mice, detected mutations that alter DNA damage response pathways. One suppressor mutation affects RBBP8, a protein that has been shown to play a role in regulating the repair of TOP2-induced DSBs (Enikanolaiye et al., 2020). This finding suggests that altering DSB pathways impacted by TOP2β dysfunction can modify Rett-like phenotypes. Thus, our findings of TOP2β regulation by MeCP2 may provide an underlying mechanism to explain these genetic findings. Given this possibility, it will be of great interest to explore the presence of TOP2β-induced DSBs in long genes of wild-type and MeCP2 mutants.

Altogether, our study has identified a molecular interaction in neurons that regulates a subset of neuronally enriched long genes. Future studies can investigate the mechanism and functional importance of this interaction and may reveal potential avenues for future therapeutic targets for MeCP2-related disorders.

## STAR Methods

### Key Resources Table

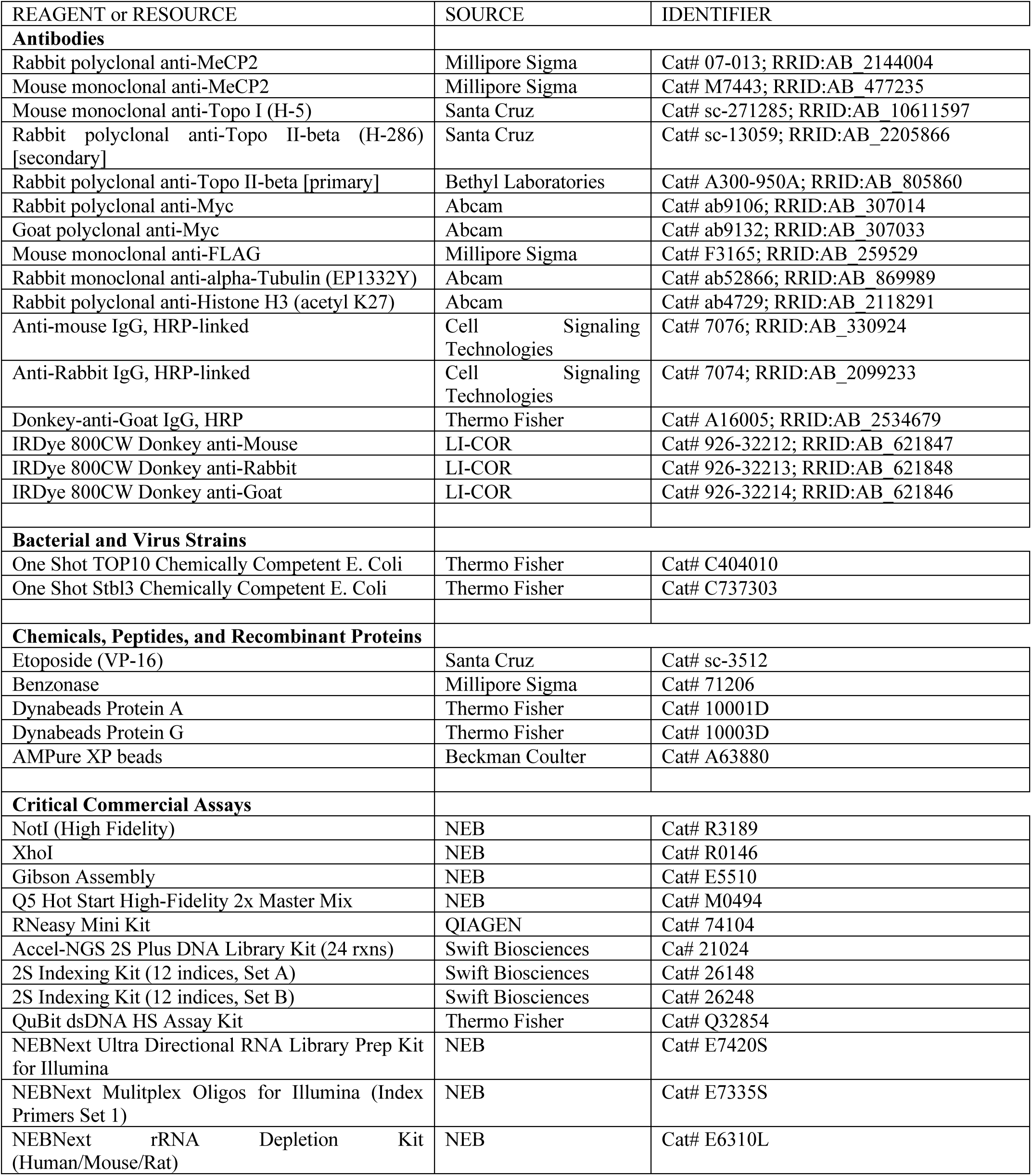
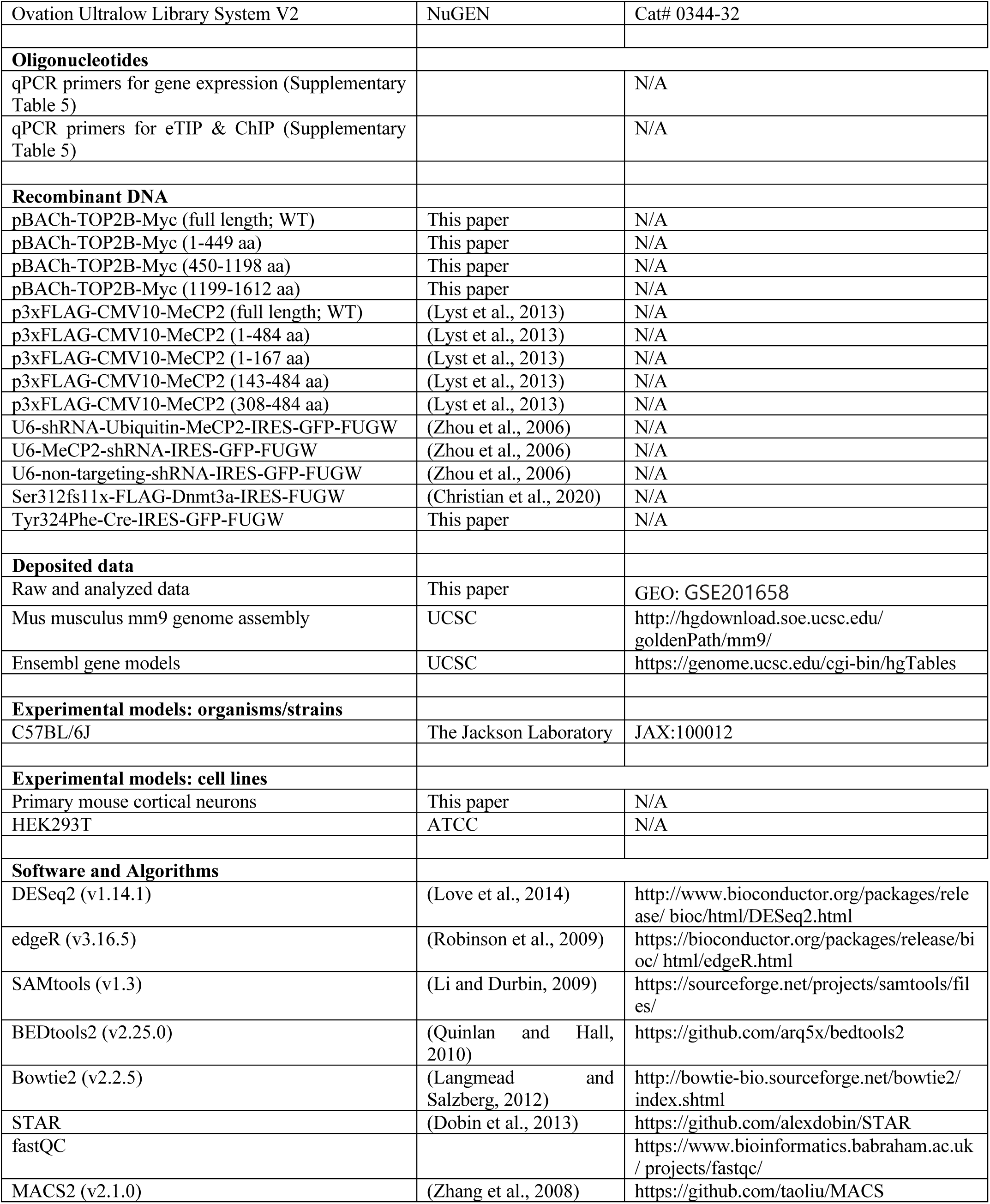
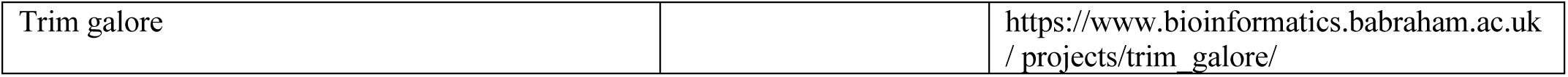

### Lead Contact and Materials Availability

Requests for reagents and resources should be directed toward the Lead Contact, Harrison Gabel.

### Experimental Model and Subject Details

#### Mouse cortical cultures

Cortical neurons were cultured from C57BL/6J E14.5 mouse embryos as described in (King et al., 2013), with some modifications. Embryonic day 14.5 (E14.5) mouse cortices were dissected in 1X DPBS, dissociated and trypsinized with TrypLE express for two 6 min incubations at 37°C, followed by DNAse treatment, to remove free-floating DNA and digest DNA from dead cells. Trituration of cells was performed with pipette to dissociate cells fully. Dissociated neurons were seeded onto 6-well plates pre-coated with poly-D-lysine (0.1mg/mL) at a density of 7.5x10^5^ cells per well (or at a density of 3x10^5^ cells per well for 12-well plates). The plates were pre-coated with poly-D-lysine (0.1mg/mL) in water, washed three times with water and washed once with Neurobasal medium before use. Neurons were cultured with neurobasal medium with 5% fetal bovine serum, GlutaMAX, B27 Supplement and Antibiotic-Antimycotic and maintained at 37°C with 5% CO_2_. Neurons were grown *in vitro* for 3 days. At DIV3 and DIV9, cells were fed with one volume of neurobasal medium supplemented with 4.84 μg/mL uridine 5’-triphosphate, 2.46 μg/mL 5-fluoro-2’-deoxyuridine, GlutaMAX, B27 Supplement and Antibiotic-Antimycotic.

### Method Details

#### Mass Spectrometry

Immunoprecipitation followed by mass spectrometry (IP-MS) was performed as described in (Mejia et al., 2013; Sowa et al., 2009), with modifications. Mouse DIV5 primary cortical neurons were infected with lentiviruses encoding FLAG-tagged baits under the CMV or neuronal Synapsin1 promoter. Lysates of DIV10 cortical neurons were subjected to immunoprecipitation using FLAG resin (Sigma), followed by 3XFLAG peptide elution (Sigma) and trichloroacetic acid (TCA) precipitation. Proteins were trypsinized (Sequencing-Grade Trypsin, Promega) and washed (3M Empore C18 media), and tryptic peptides were loaded onto an LTQ linear ion trap mass spectrometer (ThermoFinnigan). Spectra were searched against target-decoy tryptic peptide databases by CompPASS analysis.

#### Plasmids

The mouse Top2b cDNA with a C-terminal MYC tag was amplified from MGC Mouse Top2b cDNA (Dharmacon) and cloned into pBACh, a modified pCAG (cytomegalovirus enhancer fused to chicken beta-actin promoter) vector, using restriction enzyme cloning at XhoI (5’) and NotI (3’). The modified pCAG vector was generated from a pCAG-mCherry, pIRES2-EGFP (Clontech) in which the IRES and EGFP were replaced by mCherry (a gift from Dr. Jason Yi). The truncated TOP2β constructs were generated by the insertion of PCR products into pBACh empty vector using Gibson Assembly cloning (NEB). All constructs were given consensus translation initiation sequences. Human full-length MeCP2 and MeCP2 fragments were cloned into p3xFLAG-CMV (a gift from Dr. Adrian Bird) and were the same as used previously (Lyst et al., 2013). Mouse full-length Dnmt3a was the same as used previously (Christian et al., 2020). The MeCP2 shRNA and MeCP2 overexpression constructs were the same as used previously (Zhou et al., 2006).

#### Endogenous co-immunoprecipitation

Endogenous co-immunoprecipitations were carried out as described previously (Ebert et al., 2013). Forebrains from 8-week-old C57B/J mice were isolated and lysed in NP-40 lysis buffer (10 mM HEPES, pH 7.9, 3 mM MgCl2, 10 mM KCl, 10 mM NaF, 1 mM Na3VO4, 0.5 mM DTT, 0.5% NP-40, 13 complete EDTA-free protease inhibitor cocktail (Roche)), dounced 15 times with a tight pestle, and pelleted at 1,000 g. Lysates were diluted 1:1 with benzonase buffer (10 mM HEPES, pH 7.9, 3 mM MgCl2, 280 mM NaCl, 0.2 mM EDTA, 10 mM NaF, 1 mM Na3VO4, 0.5 mM DTT, 0.5% NP-40, and 13 complete EDTA-free protease inhibitor cocktail (Roche)) and digested with 250 units benzonase (Millipore) for 1h rotating at 4°C to release MeCP2 and its protein binding partners from the genome. Digested lysates were pelleted at 17,000g for 20 min at 4°C and immunoprecipitation was carried out on the supernatant using the following antibodies: MeCP2 (07-013, Millipore) or TOP2β(H-286, sc-13059, Santa Cruz), in the presence of 150mM NaCl for 2hrs while rotating at 4°C. The peptide-block control lysate was immunoprecipitated with MeCP2 antibody in the presence of a peptide to which the antibody was raised.

#### *In vitro* co-immunoprecipitation

HEK293T cells were transfected with TOP2β, MeCP2 or DNMT3A constructs using Lipofectamine 2000 (Invitrogen) and harvested after 24-48 hours. HEK293T cells were lysed in NE10 buffer (20 mM HEPES (pH 7.5), 10 mM KCl, 1 mM MgCl2, 0.1% Triton X-100 (v/v), protease inhibitors (Roche), 15 mM *β*-mercaptoethanol), dounced 15 times and pelleted 5min at 500 g. Nuclei were washed in NE10 buffer and then digested with 250 units benzonase (Millipore) for 30 min rotating at 25°C. Nuclei were resuspended in NE150 buffer (NE10 supplemented with 150mM NaCl) and incubated for 20 min. Lysates were pelleted at 16,000 g for 20 min at 4°C and supernatants were immunoprecipitated by incubating the Myc tag (ab9106, Abcam) antibody with Dynabeads Protein for 1hr at 4°C. The IP fraction was recovered by magnetic separation followed by three washes with NE10 buffer containing 150mM-300mM NaCl. The IP was then eluted from the beads with 2X NuPage LDS buffer (Invitrogen) containing *β*-mercaptoethanol.

### Immunoblotting

Nuclear extracts or IP eluates from brain tissues or HEK293T cells were resolved on 5-12% Tris-Glycine gels and transferred to nitrocellulose. Membranes were incubated overnight in the following primary antibodies: MeCP2 (Men-8, M7443, Sigma), TOP2β(H-286, sc-13059, Santa Cruz), TOP1 (H-5, sc-271285, Santa Cruz), Myc tag (ab9132, Abcam), Flag tag (F3165, Sigma), *α*-Tubulin (EP1332Y, ab52866, Abcam). Following washes, membranes were incubated with secondary antibodies conjugated to IRdye 800 and imaged with LiCOR Odyssey.

### Virus production

For lentiviral-mediated overexpression and shRNA knockdown, virus was prepared as described in (Tiscornia et al., 2006) using the MeCP2 shRNA and MeCP2 overexpression plasmids previously validated in (Zhou et al., 2006). To produce lentivirus, 10 μg of lentiviral plasmid (either shRNA-expressing or MeCP2 overexpression) was transfected into HEK293T cells along with third generating packaging plasmids pMDL (5 μg), RSV (2.5 μg), and VSVG (2.5 μg). HEK293T cells were maintained in complete DMEM media (DMEM (high glucose) media, 10% fetal bovine serum, 1% GlutaMAX, 1% penicillin/streptomycin). At 12-16 hrs following transfection, media was replaced with fresh complete DMEM media. Viruses were concentrated by ultracentrifugation 48-60 hrs after transfection and viral titers were determined by infection of HEK293T cells and were typically 0.5-1x10^5^ IFU/μL. Cultured neurons were infected on DIV3 and harvested on DIV13 for eTIP-seq, ChIP-seq or RNA-seq. In some experiments (e.g. RNA-seq of unaltered neurons) cells were transduced with control virus to control for viral infection.

### Etoposide-mediated Topoisomerase Immunoprecipitation (eTIP) protocol

eTIP experiments were performed as described in (Sano et al., 2008), with some modifications. Primary neuronal cultures from mouse cerebral cortex at 12 days in vitro (DIV12) were treated with 0.5mM etoposide (VP-16) or DMSO control in serum-free medium for 15min. The 2.1x10^6^ treated cells were lysed with 300μL of TE buffer (10mM Tris and 1mM EDTA, pH 8.0) and 1% SDS. Before sonication, 3 volumes of a buffer containing TE buffer and protease inhibitor (Complete Mini Roche) was added to the lysates. 5% of the lysates was saved for protein evaluation. To fragment DNA, the lysates were sonicated with Covaris E220 sonicator (5% Duty Factory, 140 Peak Incidence Power, 200 cycles per burst, milliTUBE 1mL AFA Fiber). Under these conditions, length of DNA fragments, as determined by TapeStation, varied from 0.5kb-1.2kb. Before immunoprecipitation (IP), 3 volumes of a buffer containing 20mM Tris-HCl (pH 8.0), 3% Triton X-100, 450mM NaCl, 3mM EDTA and protease inhibitor mixture was added to 1 volume of lysates.

The immunoprecipitation was carried out as described in (Kim et al., 2010), with some modifications. The diluted lysates were pre-cleared with 15μL of Dynabeads Protein A by rotating the tubes for 2hrs @ 4°C. After pre-clear, 3% of the lysate was saved and used as input for the IP reaction. The unbound fraction was recovered by magnetic separation. The reaction was initiated by the addition of 15μL of Dynabeads Protein A, which was pre-incubated with 2 μg of a TOP2β specific antibody. The beads suspension was incubated overnight at 4°C and the IP fraction was recovered by magnetic separation followed by two washes with low salt buffer (0.1% SDS, 1% Triton X-100, 2mM EDTA, 20mM Tris-HCl (pH 8.0), 150mM NaCl), two washes with high salt buffer (0.1% SDS, 1% Triton X-100, 2mM EDTA, 20mM Tris-HCl (pH 8.0), 500mM NaCl), two washes with LiCl buffer (0.250mM LiCl, 1% NP40, 1% deoxycholic acid (sodium salt), 1mM EDTA, 10mM Tris (pH 8.0)) and one wash with TE buffer. The IP was then eluted from the beads twice by adding elution buffer containing TE buffer and 1% SDS and incubating the samples at 65°C for 30min with brief vortexing every 10min. Elution buffer (1.5 volumes) was also added to the saved input material and this sample was processed together with the IP samples. Each eluate was treated with 10μg RNAse A and incubated for 1 hr at 37°C and then with 20mg/mL Proteinase K and incubated for 2hrs for 55°C. The IP DNA fragments were extracted with Phenol:Chloroform:Isoamyl alcohol (25:24:1, v/v). The resulting genomic DNA fragments were then purified using the QIAquick PCR purification kit (Qiagen) and DNA fragments were eluted from the columns twice.

Libraries were generated using ACCEL-NGS 2S PLUS DNA Library Kit (21024, Swift Biosciences), according to the manufacturer’s instructions and PCR amplified for 12-16 cycles. Library quality was assessed using the Agilent 4200 TapeStation (Agilent Technologies). Libraries were pooled to a final concentration of 4-10nM and 50bp reads were generated on the Illumina HiSeq 3000 with the Genome Technology Access Center, or 75bp reads were generated on the Illumina Nextseq 500 with the Center for Genome Science at Washington University in St. Louis, typically yielding 15-40 million single-end reads per sample.

### Chromatin Immunoprecipitation protocol and library preparation (ChIP-seq)

ChIP experiments were performed on 2.1x10^6^ mouse cortical neurons cultured to *in vitro* day 12 as described in (Kim et al., 2010), with some modifications. To cross-link protein-DNA complexes, media was removed from primary neurons and cross-linking buffer (20 mM HEPES-NaOH, pH 8.0, 200mM NaCl, 2 mM EDTA, 2mM EGTA) containing 1% paraformaldehyde was added for 10 minutes at room temperature. Cross-linking was quenched by adding 125 mM glycine for 5 minutes at room temperature. Cells were then rinsed 2 times in ice-cold PBS containing PMSF protease inhibitor (36978, Thermo Fisher) and collected by scrapping. Cell were lysed and nuclei isolated by incubating in L1 buffer (100 mM HEPES- NaOH, pH 7.5, 280 mM NaCl, 2 mM EDTA, 2mM EGTA, 0.5% Triton X-100, 1% NP-40, 20% Glycerol, 10 mM sodium butyrate, protease inhibitor cocktail (Roche)) for 10 minutes at 4°C. Nuclei were then pelleted by centrifugation at for 10 min at 4°C. The isolated nuclei were resuspended in L2 buffer (100mM Tris-HCl, pH 8.0, 200mM NaCl, 10 mM sodium butyrate, protease inhibitor cocktail (Roche)) and re-pelleted. The isolated nuclei were resuspended in L3 buffer (20mM Tris-HCl, pH 8.0, 2 mM EDTA, 2mM EGTA, 10 mM sodium butyrate, protease inhibitor cocktail (Roche)). Nuclei were pelleted and either stored at -80°C until use or immediately processed. Chromatin was sonicated using a Bioruptor (Diagenode) on high power mode for 50 cycles with 30 second pulses in sonication buffer (20mM Tris-HCl, pH 8.0, 2 mM EDTA, 2mM EGTA, 10 mM sodium butyrate, 0.1% Na-Deoxycholate, 0.5% SDS, protease inhibitor cocktail (Roche)).

Following sonication, the immunoprecipitation was carried out as described for eTIP and the chromatin was incubated overnight at 4°C with H3K27ac (0.025-0.1μg, Abcam 4729) and Protein A Dynabeads. H3K27ac ChIPs were performed twice from independent neuronal cultures. Following the overnight incubation, the IP was washed and eluted as described for eTIP. Libraries were generated using Ovation Ultralow Library System V2 (Tecan, 0344NB-32), according to the manufacturer’s instructions and PCR amplified for 12-16 cycles. Library quality was assessed using the Agilent 4200 TapeStation (Agilent Technologies). Libraries were pooled to a final concentration of 4-10nM and 75bp reads were generated on the Illumina Nextseq 500 with the Center for Genome Science at Washington University in St. Louis, typically yielding 15-40 million single-end reads per sample.

### Total RNA Isolation and library preparation (RNA-seq)

Neuronal cultures dissected and transduced (with MeCP2 KD, MeCP2 OE or control virus) on the same days, constituted technical replicates, whereas, Neuronal cultures dissected and transduced on independent days, constituted biological replicates. Cultured neurons were harvested directly using RLT buffer and homogenized in the QIAshredder spin column (Qiagen). To isolate RNA, the RNeasy Mini Kit (Qiagen) according to the manufacturer’s instructions. RNA libraries were generated from 250ng total RNA with rRNA depletion (NEBNext, E6310) and NEBNext Ultra Directional RNA Library Prep Kit for Illumina (NEBNext, E7420), using a modified amplification protocol (37°C, 15 minutes; 98°C, 30 s; (98°C, 10 seconds; 65°C, 30 seconds; 72°C, 30 seconds) x 13 cycles; 72°C, 5 minutes; 4°C, hold). RNA libraries were pooled at a final concentration of 8-10nM and single end 75bp reads were generated on the Illumina Nextseq 500 with the Center for Genome Science at Washington University in St. Louis, typically yielding 20-40 million single-end reads per sample.

### Quantification and Statistical Analysis

#### Etoposide-mediated Topoisomerase Immunoprecipitation sequencing analysis

eTIP-seq was analyzed as previously described for ChIP-seq data in (Clemens et al., 2020). Sequenced reads were mapped to the mm9 genome using bowtie2 alignment and reads were extended based on library sizes and deduplicated to consolidate PCR duplicate reads. Deduplicated reads were used to quantify read density normalized by the number of reads per sample and by read length in basepairs. Bedtools coverage -counts parameter was used to quantify eTIP signal and ChIP signal.

For eTIP, the signal was quantified at promoter associated region, defined as 1kb downstream to 3kb downstream of the TSS, and the gene body, defined as 3kb downstream of the TSS to the end of the transcript, based on our Ensembl gene models. edgeR was then used to determine differential eTIP-signal across conditions. Data were visualized using UCSC genome browser (http://genome.ucsc.edu).

Aggregate plots of eTIP signal at genes and enhancers were generated by calculating eTIP/Input for equally sized bins for the specified windows using Bedtools coverage -hist parameter and custom R and python scripts. For aggregate analysis at genes, the genes less than 3kb in length were filtered out in order to capture the signal at the promoter associated regions without conflating the ends of genes with this region. In some instances, the genes are further filtered such that the lengths of the genes plotted is equal to or greater than the aggregate length being plotted (Clemens et al., 2020) .

Resampling control analysis was performed with custom scripts, by moving true TOP2β peak randomly around the genome with the criteria that the shuffled peaks do not overlap each other however could at random overlap a true TOP2β peak.

#### Chromatin Immunoprecipitation sequencing analysis

ChIP-seq analysis was performed as previously described (Clemens et al., 2020). Sequenced reads were mapped to the mm9 genome using bowtie2 alignment and reads were extended based on library sizes and deduplicated to consolidate PCR duplicate reads. Deduplicated reads were used to quantify read density normalized by the number of reads per sample and by read length in basepairs. Bedtools coverage -counts parameter was used to quantify ChIP signal.

For ChIP, the signal was quantified at the TSS and enhancers. Enhancers in this study were defined by requiring the presence of H3K27ac peaks that occur outside of a known TSS region (TSS +/-500bp). H3K27ac peaks were identified using MACS2 peak calling algorithm, in which the ChIP input was used as background signal, using the following parameters: macs2 call peak --nomodal -q 0.05. Bedtools intersect was used to identify H3K27ac peaks that did not overlap with gene promoter regions. These filtered H3K27ac peaks were defined as enhancers. As noted, this may have led to the exclusion of some subthreshold regions of H3K27ac enrichment that may represent true regulatory elements.

#### RNA sequencing analysis

RNA sequencing analysis was performed as previously described in (Clemens et al., 2020). Raw FASTQ files were trimmed with Trim Galore, using a quality filter of 20, followed by filtering out rRNA sequences using Bowtie2. Remaining reads were aligned to mm9 using STAR (Dobin et al., 2013) with default parameters. Reads mapping to multiple regions in the genome were then filtered out, and uniquely mapped reads were converted to BED files and separated into intronic and exonic reads. Finally, reads were assigned to genes using bedtools coverage -counts parameter (Quinlan and Hall, 2010).

For gene annotations, we defined a “flattened” list of the longest transcript for each gene, generated on Ensgene annotations and obtained from the UCSC table browser. For each gene, Ensembl IDs were matched up to the MGI gene names. Then, for each unique MGI gene name, the most upstream Ensgene TSS and the most downstream TES were taken as the gene’s start and stop. Based on these Ensembl gene models, we defined TSS regions, TSS-adjacent regions and gene bodies. DESeq2 was run using default parameters on exonic reads from MeCP2 KD and MeCP2 OE cultured neurons (n = 3 per condition), to identify differentially expressed genes.

### Data and Software Availability

The accession number for the raw and processed data in this paper is Gene Expression Omnibus, GEO: GSE201658.

## Acknowledgements

We thank Dennis Wu for guidance and support on genomic and bioinformatic analysis. We thank members of the Gabel lab, as well as J. Yi, J. Dougherty, A. Yoo, T. Turner, G. Zhao and T. Li for helpful discussions and critical reading of the manuscript. We thank J. Hoisington-Lopez and M. Crosby at the Center for Genome Sciences (CGS) and the Genome Technology Access Center (GTAC) at Washington University in St. Louis for sequencing services. This work was supported by Klingenstein-Simons Fellowship Fund, the G. Harold and Leila Y. Mathers Foundation, the Brain and Behavior Research Foundation, and the Simons Foundation Autism Research Initiative; and NIMH R01MH117405, NINDS R01 NS04102 to H.W.G.

## Author Contributions

Conceptualization, S.A.N. and H.W.G.; Methodology, S.A.N. and H.W.G.; Formal analysis, S.A.N.; Investigation: designed, executed and analyzed all experiments, S.A.N. Performed IP-MS, Y.I. Assisted with cloning and lentivirus production, C.I.A. Performed endogenous co-IP, H.W.G.; Writing – original draft, S.A.N. and H.W.G.; Writing – reviewing & editing, all of the authors; Supervision, H.W.G., A.B; Funding acquisition, H.W.G., A.B.

## Supplemental Information

Supplemental Information includes five tables and two figures.

**Table S1.** Mass spectrometry Table; Related to Figure 1

**Table S2.** Number of reads per sequencing sample

**Table S3.** Gene Expression Table; Related to Figure 3, 4, S1, S2

**Table S4.** TOP2βeTIP-seq & H3K27ac ChIP-seq Table; Related to Figure 3, 4, S1, S2

**Table S5.** qPCR Primer Sequences for Gene Expression, TOP2β eTIP, H3K27ac ChIP; Related to Figure 3, 4, S1, S2

---

**Supplementary Figure 1.**
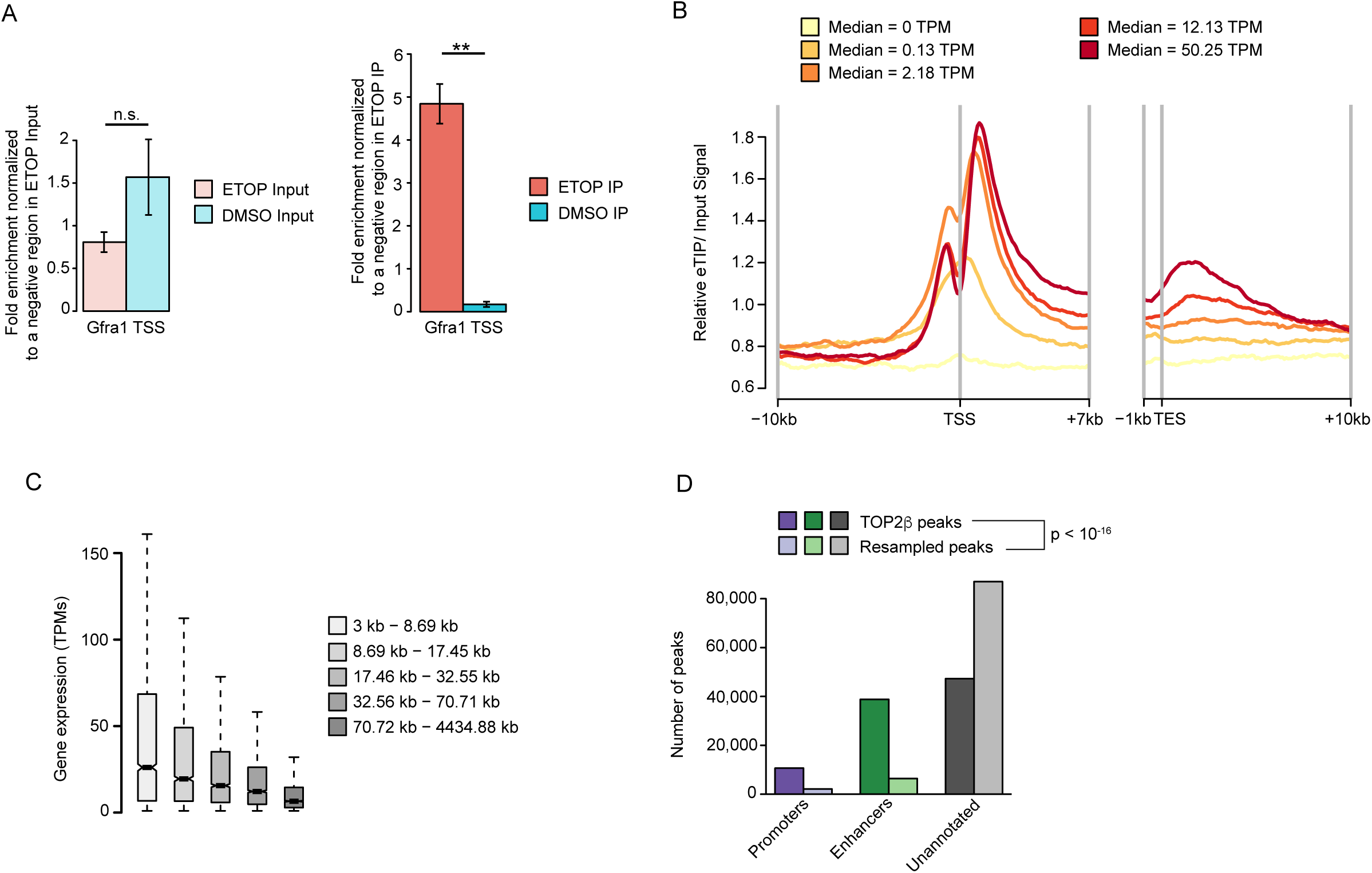
**Analysis of TOP2β activity in neurons by eTIP-seq.** (A) Quantitative PCR (qPCR) analysis of input and IP DNA isolated from ETOP-treated vs. DMSO-treated DIV12 cultured cortical neurons. Data were normalized to a negative control intergenic region in the ETOP-treated condition (see *Methods*). Data are mean ± SEM for six (input) and seven (IP) independent experiments. Two-tailed unpaired t-test (n.s., not significant; **, p < 10^-5^). (B) Aggregate plot of input-normalized eTIP signals at genes divided into quantiles of gene expression. The median gene expression (TPM) for each group is indicated. Average signal around the transcription start site (TSS) and transcription end site (TES) is shown. (C) Boxplot of exonic RNA expression from DIV12 wild-type cultured cortical neurons for genes divided in quantiles of gene length. (D) Bar plots of the genomic distribution of TOP2β peaks and resampled TOP2βpeaks at promoters, enhancers and unannotated regions. Resampling and statistical analysis was performed as described in **Figure 3F**.

**Supplementary Figure 2.**
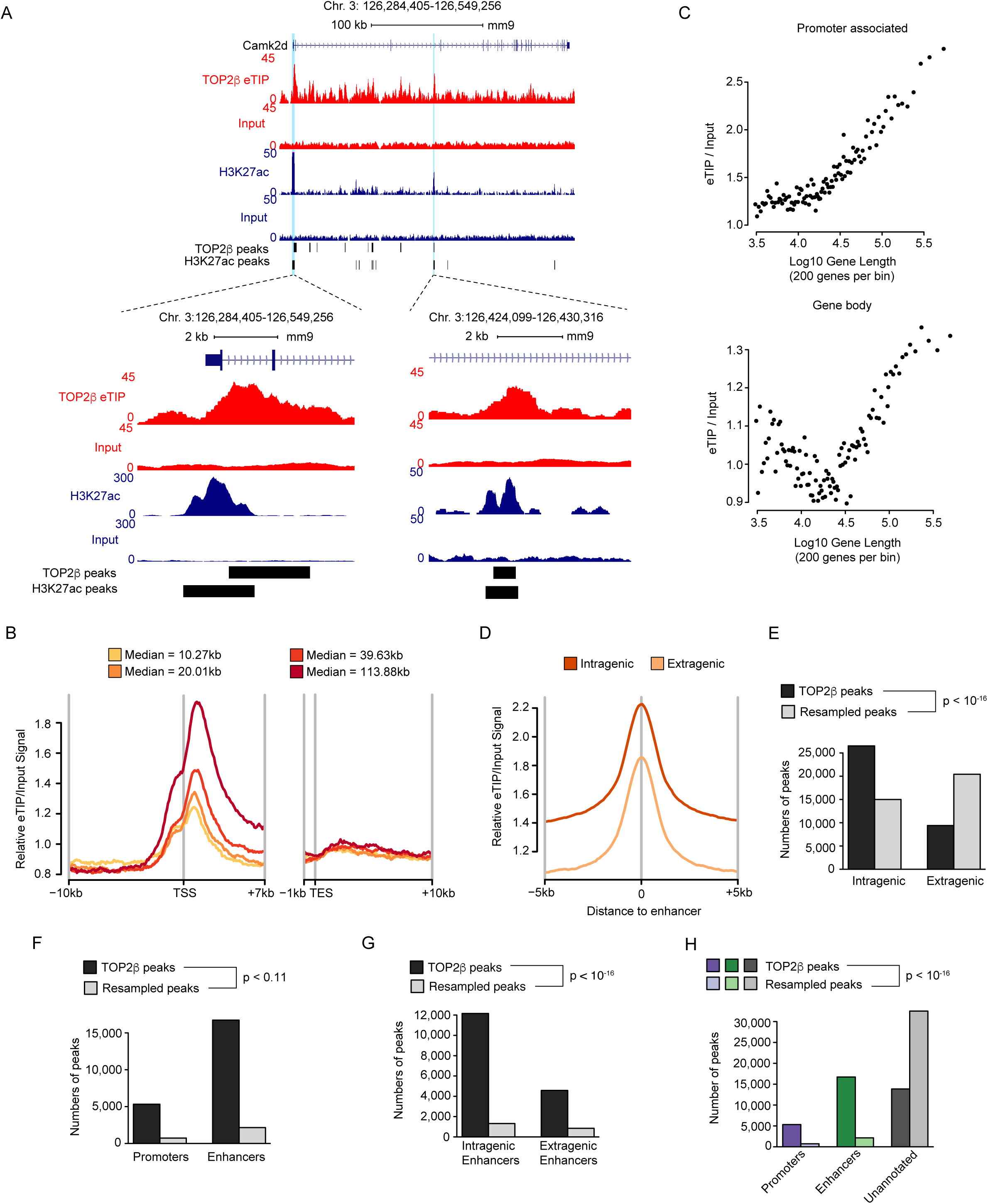
**Validation of TOP2β activity profiles in neurons by eTIP-seq with a second TOP2β antibody.** (A) Genome browser snapshots of eTIP-seq and H3K27ac ChIP-seq from DIV12 cultured cortical neurons at an example long gene, *Camk2d*. Regions corresponding to H3K27ac peaks (putative enhancers and promoters) are highlighted in blue. Bottom left, zoomed in view of the promoter region of the *Camk2d* gene. Bottom right, zoomed in view of an enhancer region of the *Camk2d* gene. MACS2 called-peaks of TOP2β eTIP and H3K27ac are indicated below the tracks. Gene annotation and scale are depicted above. (B) Aggregate plot of input-normalized eTIP signals at genes divided into quantiles of gene length. The median gene length for each group is indicated. Average signal around the transcription start site (TSS) and transcription end site (TES) is shown. Mean values plotted for 100bp bins. (C) Running average plots of input-normalized eTIP signal at the promoter associated region (top) and gene bodies (bottom), from eTIP-seq. Averages are shown for bins of 200 genes ranked according to gene length. (D) Aggregate plot of input-normalized eTIP signal at putative intragenic and extragenic enhancers genome-wide in cultured cortical neurons. Putative enhancers were defined by H3K27ac peaks that did not overlap the TSS of a gene. The plot is centered at the midpoint for each enhancer and the surrounding 10kb region. Mean values plotted for 100bp bins. (E) Bar plots of the genomic distribution of TOP2β peaks and resampled TOP2βpeaks at intragenic and extragenic regions. Resampling and statistical analysis were performed as described in **Figure 3F**. (F) Bar plots of the genomic distribution of TOP2β peaks and resampled TOP2βpeaks at promoters and enhancers. Resampling and statistical analysis were performed as described in **Figure 3F**. (G) Bar plots of the genomic distribution of TOP2β peaks and resampled TOP2βpeaks at intragenic enhancers and extragenic enhancers. Resampling and statistical analysis were performed as described in **Figure 3F**.

